# PF-D-Trimer, a broadly protective SARS-CoV-2 subunit vaccine: immunogenicity and application

**DOI:** 10.1101/2022.09.26.509414

**Authors:** Zhihao Zhang, Jinhu Zhou, Peng Ni, Bing Hu, Shuang Deng, Qian Xiao, Qian He, Gai Li, Yan Xia, Mei Liu, Cong Wang, Zhizheng Fang, Nan Xia, Zhe-Rui Zhang, Bo Zhang, Kun Cai, Normand Jolicoeur, Yan Xu, Binlei Liu

## Abstract

The COVID-19 pandemic, caused by the SARS-CoV-2 virus, has had and still has a considerable impact on global public health. One of the characteristics of SARS-CoV-2 is a surface homotrimeric spike protein, the primary responsible for the host immune response upon infection. Here we show the preclinical studies of a broad protective SARS-CoV-2 subunit vaccine developed from our Trimer Domain platform using the Delta spike protein, from antigen design to purification, vaccine evaluation and manufacturability. The prefusion trimerized Delta spike protein, PF-D-Trimer, was highly expressed in Chinese hamster ovary (CHO) cells, purified by a rapid one-step anti-Trimer Domain monoclonal antibody immunoaffinity process and prepared as a vaccine formulation with an adjuvant. The immunogenicity studies demonstrated that this vaccine candidate induces robust immune responses in mouse, rat and Syrian hamster models. It also protects K18-hACE2 transgenic mice in a homologous virus challenge. The neutralizing antibodies induced by this vaccine display a cross-reactive capacity against the ancestral WA1 and Delta variants as well as different Omicron, including BA.5.2. The Trimer Domain platform was proven to be a key technology in the rapid production of the PF-D-Trimer vaccine and may be crucial to accelerate the development of updated versions of SARS-CoV-2 vaccines.

## Introduction

The appearance of the SARS-CoV-2 virus in 2019 quickly became a major public health problem with the rapid spread of the COVID-19 pandemic knowing no borders ^1^. Like most enveloped RNA viruses, SARS-CoV-2 uses a trimeric surface protein, the spike (S) protein, when infecting a host cell. S protein is responsible for the attachment by binding to a cellular receptor, hACE2 ^2^ and then mediates viral entry by membrane fusion ^3^. Several research results and available vaccines have already demonstrated the importance of the spike protein as an ideal target for vaccine development ^4^. Current SARS-CoV-2 vaccines can be categorized into four different classes, summarily: nucleic acids, RNA or DNA, which encode part of the genetic information of the virus; inactivated vaccines which consist of a virus being physically treated to render it incapable of producing disease; viral vector vaccines, for example adenovirus with limited replication capacity, which encodes part of the SARS-CoV-2 genome to introduce it into a host cell; recombinant protein subunit vaccines, which do not use viral genetic material, but rather full-length viral proteins or fragments thereof, either packaged or not in nanoparticles for better delivery and uptake by cells responsible for immunity ^5,6^. It has been known for several years that major epitopes of the S protein of coronaviruses only exist in its trimeric form, and are so trimer restricted ^7^. Recently published research results also support the concept that the trimeric form of the S protein adopts a conformation containing important vaccine epitopes ^8^. As demonstrated in a recent publication, the Delta variant has the potential of becoming far more problematic than Omicron ^9^ and low titers of neutralizing antibodies are associated with SARS-CoV-2 Delta breakthrough infections in vaccinated patients ^10^. There is also evidence for SARS-CoV-2 Delta and Omicron co-infections and recombination ^11,12^. Sera from unvaccinated or vaccinated Delta-wave intensive care unit (ICU) patients strongly neutralize Omicron BA.4/5 and BA.2.12.1 ^13^.

PF-D-Trimer, is a subunit SARS-CoV-2 vaccine candidate, consisting of the recombinant S-glycoprotein from the Delta variant in its prefusion form ^14^ and trimerized by fusion with our proprietary Trimer Domain (TD) ^15^. We demonstrate that PF-D-Trimer adjuvanted with alum and CpG 1018 induces a strong immune response in the form of circulating and neutralizing antibodies against the original WA1 virus, as well Delta and different Omicron variants, including BA.2.2 and BA.5.2. The formulation also induces a Th1-biased cellular immune responses in animal models and protect K18-hACE2 H11 transgenic mice in a homologous challenge study. These results support our vaccine strategy of using the Delta variant S protein as an antigen. In addition, we describe here the use of our TD platform which not only allows the stabilization of the S protein in a trimeric form, but also its simplified and rapid one-step purification by immunoaffinity, allowing the development of a streamlined chemistry manufacturing and control (CMC) strategy.

## Results

### High level expression, purification and characterization of PF-D-Trimer from bioreactor cultures

PF-D-Trimer mimics the native trimeric prefusion structure of the S protein, as found on the virus surface, and consists of the complete ectodomain of the S protein of the SARS-CoV-2 Delta variant fused C-terminally to the TD trimerization domain (Fig. 1A). A high-level production process was developed following the transfection of CHO cells with an expression vector containing a CHO codon-optimized cDNA encoding PF-D-Trimer, the selection of the best performing clones and process optimizations. This process makes it possible to achieve production levels of more than 500 mg/L of PF-D-Trimer as a secreted protein in the culture medium. The expression process is robust as demonstrated in the side-by-side analysis of two different batches by SDS-PAGE which yield nearly identical expression levels (Fig. 1B). Reducing and non-reducing SDS-PAGE analysis demonstrated that the purified PF-D-Trimer is a self-associating homotrimer stabilized by interchain disulfide bonds (Fig. 1C). Side-by-side analysis of two purification batches of PF-D-Trimer demonstrates the reproducibility of the purification process, the two batches having similar apparent levels of purity after silver staining (Fig. 1C). Under reducing conditions PF-D-Trimer appears as a single form with a molecular weight of around 170 kDa. Under non-reducing conditions, PF-D-Trimer appears as a single high molecular weight form, thus demonstrating that the protein is not cleaved by proteases produced by CHO cells (Fig. 1C). The PF-D-Trimer eluted from the one-step α−TD trimerization domain monoclonal antibody immunoaffinity chromatography was assessed by SEC-HPLC, showing a >99% main peak around 700 kDa, without obvious aggregation or fragmentation (Fig. 1D). Data from electron microscopy confirm that immunoaffinity-purified PF-D-Trimer sample contains assembled spike trimers (Fig. 1E).

**Fig. 1.**
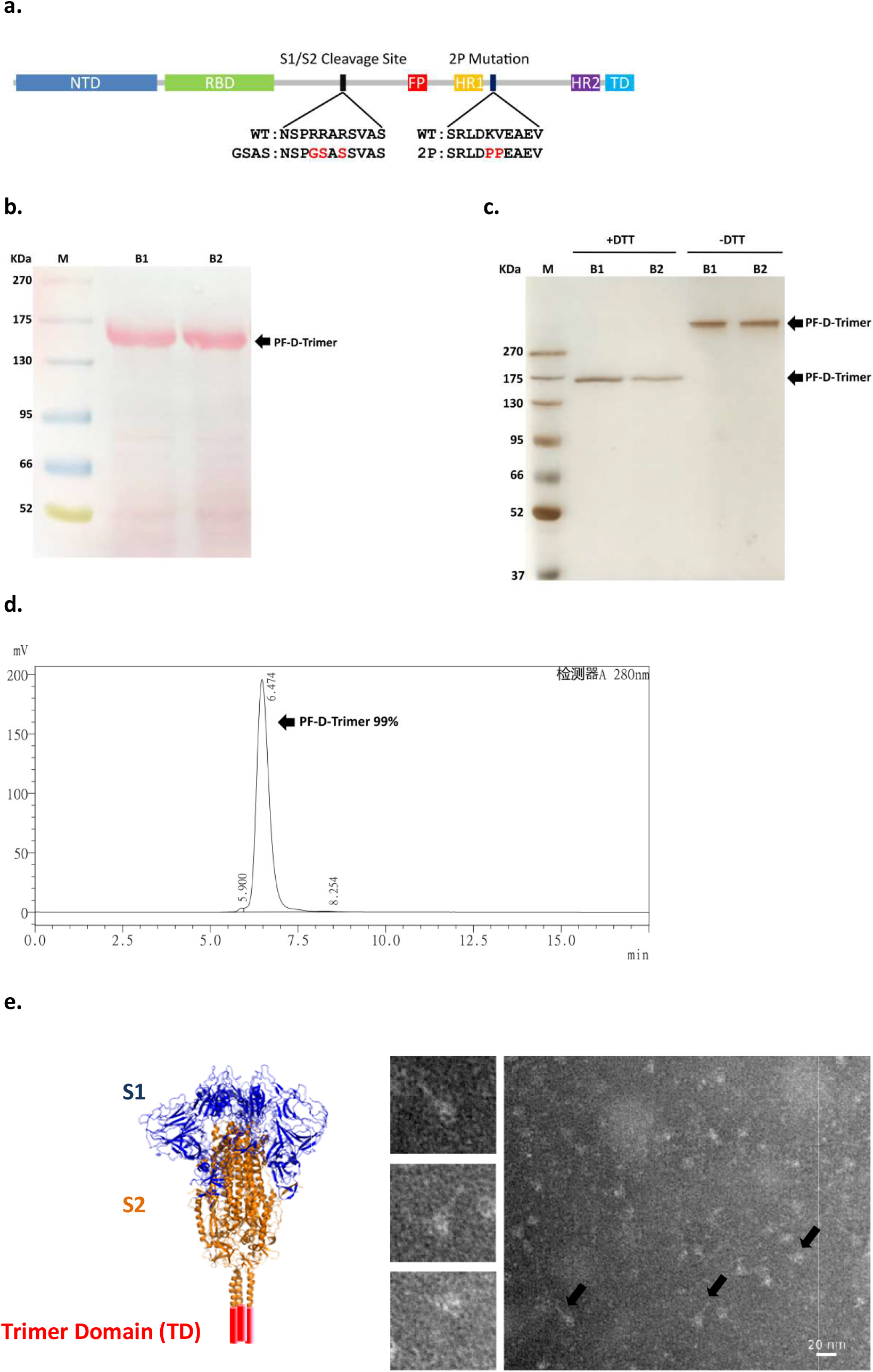
Expression, Purification and Characterization of PF-D-Trimer. **(a)** Schematic representation of the S ectodomain of Delta variant SARS-CoV-2 Trimer Domain fusion protein (PF-D-Trimer). **(b)** Analysis by reducing SDS-PAGE with Ponceau S staining of high-level expression of PF-D-Trimer from CHO cells in bioreactor cultures over a period of 11 days. Samples from two batches (B1, B2) were analyzed side by side. Each well contains 16 µL of CHO cells culture supernatant. **(c)** PF-D-Trimer is a disulfide bond-linked homotrimeric protein as analyzed by SDS-PAGE with silver staining under reducing (+DTT) and non-reducing (-DTT) conditions. Samples from two purification batches (B1, B2) were analyzed side by side. Each well contains 1 µg of PF-D-Trimer as quantified by molar absortivity at 280 nm. **(d)** SEC-HPLC analysis of the immunoaffinity purified PF-D-Trimer showing a MW of approximately 700 KDa and a purity of more than 99%. **(e)** PF-D-Trimer schematic diagram and representative negative-stain EM images with homotrimeric S protein in the prefusion conformation. The images on the left are partial enlargements of the image on the right as indicated with the black arrows. The sample contains assembled spike trimers. Scale bar 20 nm.

### Induction of immune response by alum plus CpG 1018 adjuvanted PF-D-Trimer in mice

The immunogenicity of alum and CpG 1018 adjuvanted PF-D-Trimer as vaccine was first evaluated in C57BL/6 mice. Sera from these mice were evaluated for the amount of anti-spike IgG. Sera from the PBS formulation injected mice only showed titers at background level (result not shown). At Day 21 (Fig. 2A), the antibody reciprocal GMT titers of the 5 μg, 10 μg and 20 μg groups reached 6.5×10^5^, 4.6×10^5^ and 1.2×10^6^, respectively. The 10 μg and 20 μg groups displayed significant differences in antibody titers (p=0.0329). At Day 35 (Fig. 2B), the antibody reciprocal GMT titers of the 5 μg, 10 μg and 20 μg groups reached 9.3×10^5^, 1.1×10^6^ and 1.2×10^6^, respectively, and there was no significant difference in the antibody titers among the groups. Together these results demonstrate that the antibody responses were increased by the booster immunization.

**Fig. 2.**
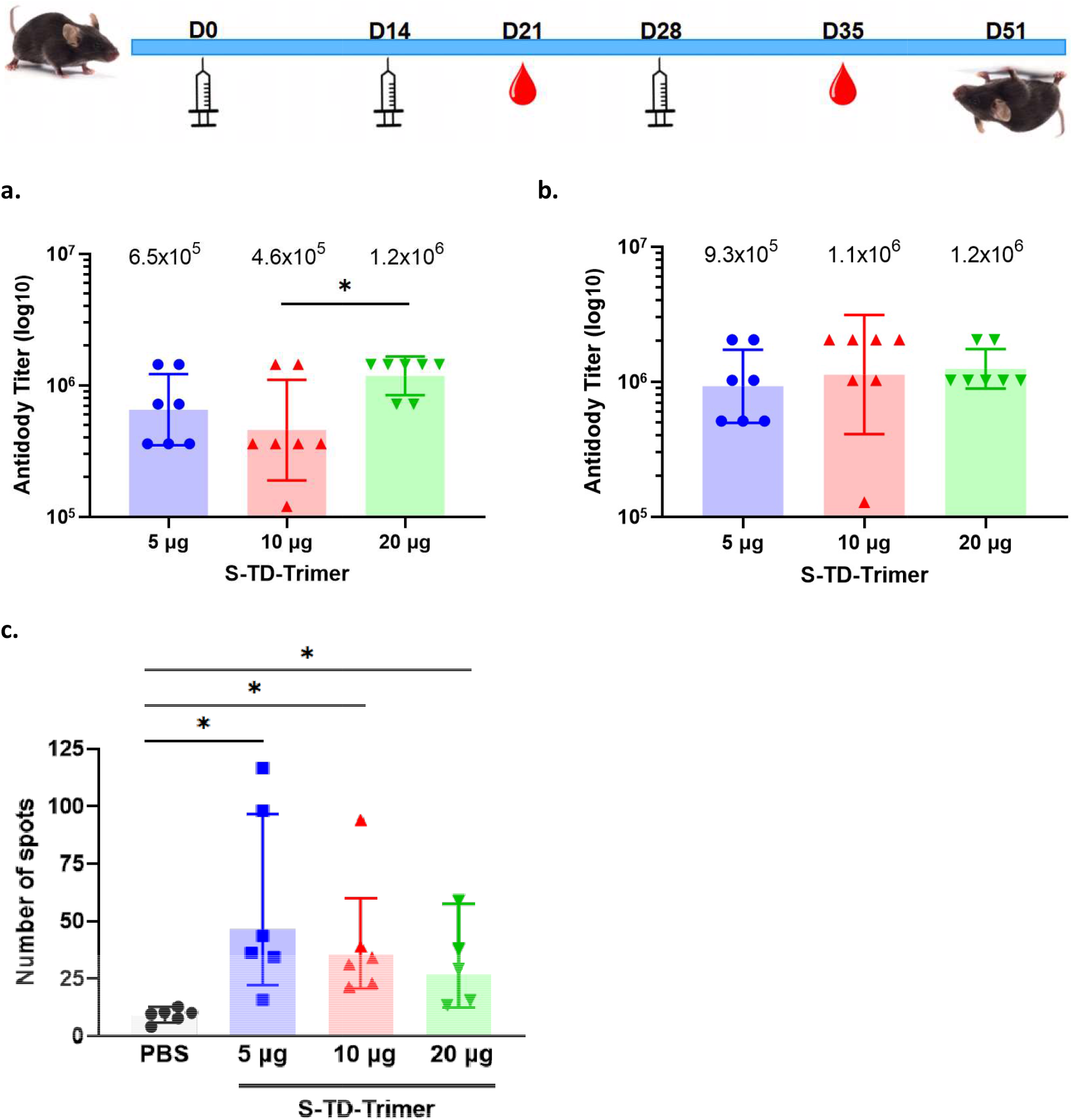
Immunogenicity of PF-D-Trimer in mice. C57BL/6 mice were immunized with three different doses of PF-D-Trimer formulated with alum plus CpG 1018, as indicated in the top panel and described in Material and Methods. Sera were collected and the immune responses were analyzed 2 weeks after the second (Day 21) and the third (Day 35) injections. **(a)** PF-D-Trimer binding antibody ELISA titers at Day 21. **(b)** PF-D-Trimer binding antibody ELISA titers at Day 35. **(c)** Detection of Th1 (IFN-γ) by ELISpot after stimulation by S protein. Points represent individual animal, bars indicate geometric mean titers (GMT) responses (with 95% confidence intervals [CI]) for antibody assays and geometric mean counts (GMC) responses (with 95% CI) for ELISpot. * p≤0.05, *** p≤0.001, ****p≤0.0001. • 5 µg group, ▴10 µg group, ▾ 20 µg group.

Mice immune sera were further tested to identify whether the vaccine-adjuvant system could induce Th1 responses in the vaccinated mice. Th1 IFNγ responses were measured in the splenocytes of the immunization groups (Fig. 2C). Compared with the PBS control group, the 5 μg, 10 μg and 20 μg groups had significant differences in the number of anti S protein specific IFNγ producing T cells. These results demonstrate that the formulated PF-D-Trimer is promoting the development and activation of Th1 cells in C57BL/6 mice.

### Induction of neutralizing antibodies by alum plus CpG 1018 adjuvanted PF-D-Trimer in Sprague-Dawley rats

We also examined the immunogenicity of alum and CpG 1018 adjuvanted PF-D-Trimer in Sprague-Dawley rats. During the progression of the COVID-19 pandemic many variants of SARS-CoV-2 emerged. Among them is Omicron, a highly mutated variant of concern highly resistant to vaccine-induced antibody neutralization ^16^. Thereby, the immune sera were tested for their neutralization capabilities against the Delta and Omicron BA.2.2 and BA5.2 variants of SARS-CoV-2. The reciprocal ID_50_ GMT of 10 µg PF-D-Trimer adjuvanted with both alum and CpG 1018 reached 21478 for Delta, 1600 for Omicron BA.2.2 and 2344 for Omicron BA.5.2 after three doses at Day 71 (Fig. 3A). The reciprocal ID_50_ GMT of 30 µg adjuvanted PF-D-Trimer reached 28076 for Delta, 2042 for Omicron BA.2.2 and 4519 for Omicron BA.5.2 after three doses at Day 71 (Fig. 3B). Thus, neutralizing antibodies were generated against the three variants using the adjuvanted PF-D-Trimer, the highest neutralization being with Delta (homologous immunization), followed by Omicron BA.5.2 and then Omicron BA.2.2. Neutralization titers for Omicron BA.5.2 were higher than BA.2.2, regardless of the dose.

**Fig. 3.**
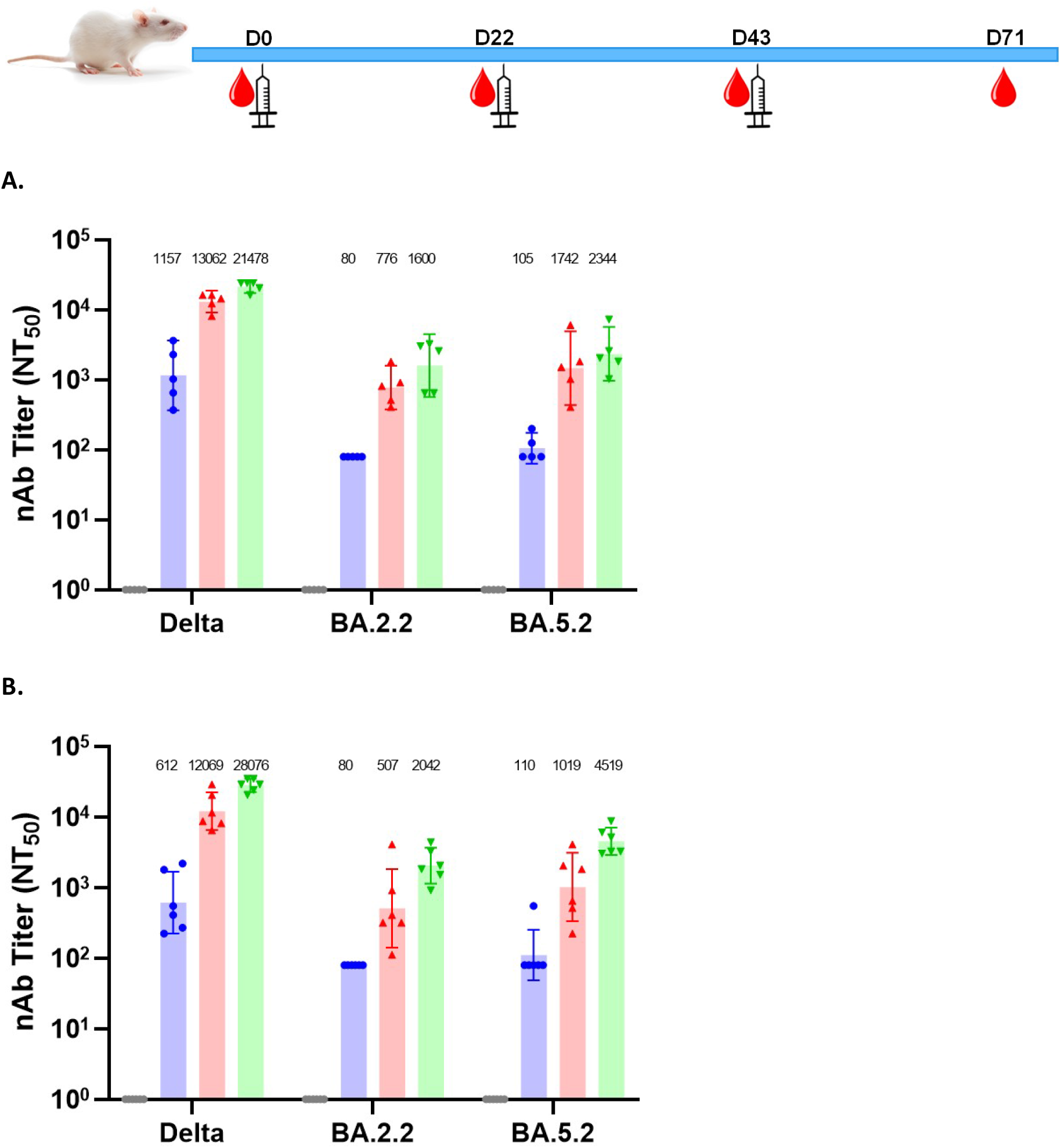
Induction of neutralization antibodies by PF-D-Trimer in Sprague-Dawley rats. Sprague-Dawley rats were immunized with two different doses of PF-D-Trimer formulated with alum plus CpG 1018 as indicated in the top panel and as described in Material and Methods. Sera were collected and the neutralization activities against Delta and Omicron BA.2.2 and BA5.2 were analyzed as described in Material and Methods. **(a)** Sera neutralization activities with 10 µg of adjuvanted PF-D-Trimer. **(b)** Sera neutralization activities with 30 µg of adjuvanted PF-D-Trimer. Points represent individual animal, bars indicate geometric mean titers (GMT) responses (with 95% confidence intervals [CI]). • Day 22, ▴Day 43, ▾ Day 71.

### Induction of immune response by alum plus CpG 1018 adjuvanted PF-D-Trimer in Syrian golden hamsters

We evaluated the immunogenicity and potential efficacy of PF-D-Trimer formulated with or without alum and with or without CpG 1018 after two or three injections containing 5 µg or 10 µg of PF-D-Trimer in Syrian golden hamsters. Twenty days after the second injection (Fig. 4A), all the immunization groups including the non-adjuvanted protein group were showing a humoral immune response. The geometric means were 8.9×10^5^ and 3.6×10^6^ for the 5 µg groups adjuvanted with alum or alum plus CpG 1018 respectively, the later giving a significantly higher GMT titer. In the 10 µg groups the GMT were 1.1×10^6^ and 2.8×10^6^ for the formulations adjuvanted with alum or alum plus CpG 1018 respectively, the later being significantly higher. Interestingly, the unadjuvanted PF-D-Trimer was still giving a GMT of 2.9×10^5^ even though significantly lower than the adjuvanted formulations. The third injection boosted the GMT for the 5 µg (results not shown) and the 10 µg groups (Fig. 4B), although these did not differ significantly from the second injection, regardless of whether the formulations were adjuvanted or not in each group.

**Fig. 4.**
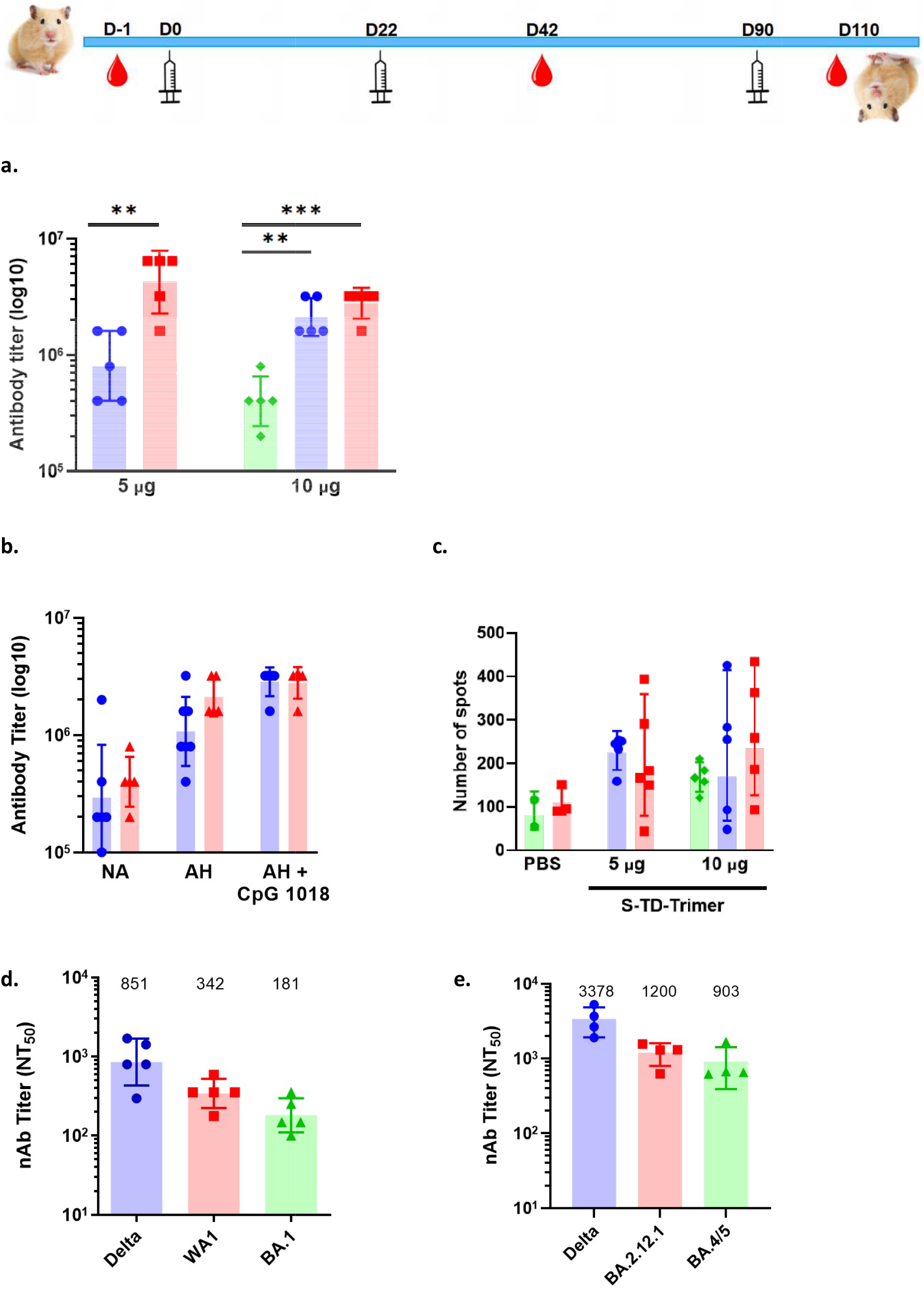
Immunogenicity of PF-D-Trimer in Syrian golden hamsters. Syrian golden hamsters were immunized with two different doses of PF-D-Trimer formulated or not with alum plus CpG 1018, as indicated in the top panel and as described in Material and Methods. Sera were collected and the immune responses were analyzed twenty days after the second and third injections (Day 42 and Day 110). **(a)** PF-D-Trimer binding antibody ELISA titers at Day 42. ◆ Not Adjuvanted, • Alum, ▪ Alum plus CpG 1018. **(b)** Comparison of different PF-D-Trimer formulations binding antibody ELISA titers at Day 42 and Day 110. NA, not adjuvanted; AH, alum; AH + CpG 1018, alum plus CpG 1018. • Second immunization, ▴Third Immunization. **(c)** Detection of Th1 (IFN-γ) by ELISpot after stimulation by S protein. ◆ Not Adjuvanted, • Aluminum Hydroxide, ▪ Alum plus CpG 1018. **(d)** Sera neutralization activities against WA1 and the Delta and Omicron BA.1 variants. **(e)** Sera neutralization activities against SARS-CoV-2 Delta, BA.2.12.1 and BA.4/5 pseudotyped viruses. Points represent individual animal, bars indicate geometric mean titers (GMT) responses (with 95% confidence intervals [CI]) for antibody assays and geometric mean counts (GMC) responses (with 95% CI) for ELISpot. * *p≤0.01, *** p≤0.001.

Splenocytes from hamster groups harvested after immunization were tested for their Th1 IFNγ response (Fig. 4C). There was no significant difference in the numbers of anti PF-D-Trimer-specific IFNγ producing T cells between the 5 µg and 10 µg groups irrespectively of the formulations. Moreover, the protein alone is also able to generate a T cell response. Taken together, these results suggest that PF-D-Trimer direct the hamsters’ T cells towards a Th1 phenotype.

Sera from hamsters that had received three 5 µg injections of PF-D-Trimer formulated with alum plus CpG 1018 were used to assess their neutralization activities toward the ancestral WA1 and the Delta strains as well as Omicron variants of SARS-CoV-2 (Fig. 4D). The reciprocal ID_50_ GMT were 851 for Delta, 342 for WA1 and 181 for Omicron BA.1. Thus, neutralizing antibodies were generated against the three strains.

The sera were also used to assess their neutralization activities toward SARS-CoV-2 Delta, BA.2.12.1 and BA.4/5 pseudotyped viruses. The reciprocal ID_50_ GMT were 3378 for Delta, 1200 for BA.2.12.1 and 903 for BA.4/5 (Fig. 4E). Similar to the results of the experiments carried out with the real viruses, the results with the pseudotyped viruses also show a neutralization activity against the three strains.

### Immunogenicity and protective efficacy of adjuvanted PF-D-Trimer in K18-hACE2 H11 transgenic mice

The protective efficacy of PF-D-Trimer adjuvanted or not with alum plus CpG 1018 was examined in K18-hACE2 mice. hACE2 mice develop respiratory disease resembling severe COVID-19 and is a suitable model to study pathogenesis and immune responses ^17^.

We examined the immunogenicity of alum plus CpG 1018 adjuvanted PF-D-Trimer in K18-hACE2 mice. The immune sera collected after two immunizations were evaluated for the amount of anti-spike IgG (Fig. 5A). The antibody reciprocal GMT of the 30 μg group reached 1.4×10^7^ while for the 10 μg group, although the titer was lower compared to the 30 μg group, it was still significantly higher than the control groups with a titer of 2.7×10^6^. Mice vaccinated with only the adjuvant and intranasally infected with the Delta strain (Fig. 5B) were showing a decrease in weight and died at 4 or 5 DPI (Fig. 5C). For the mice vaccinated with 10 μg or 30 μg of adjuvanted PF-D-Trimer, the body weight slightly decreased in the first 2 days after infection, and subsequently recovered. The mice in the PF-D-Trimer vaccinated groups were in good condition at 7 DPI and all survived. The viral load in the mice’s lung tissue was detected on the third day after infection (Fig. 5D). The live virus titer in lung tissue was below the minimum detection limit (100 PFU/g) in the mice from the 10 μg and 30 μg of adjuvanted PF-D-Trimer groups, while load from the mice of the adjuvant only group reached more than 1 × 10^5^ PFU/g of lung tissue.

**Fig. 5.**
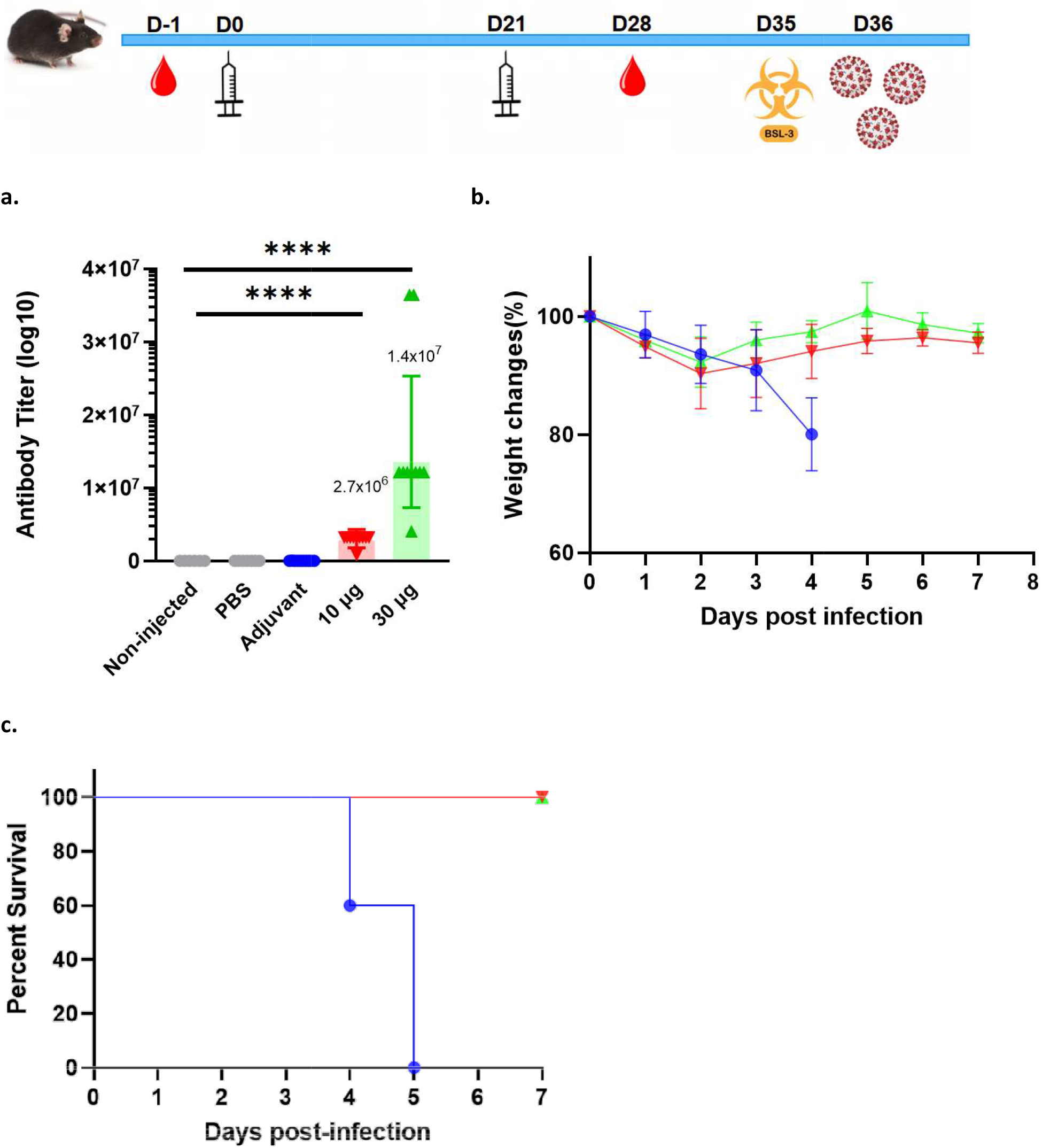
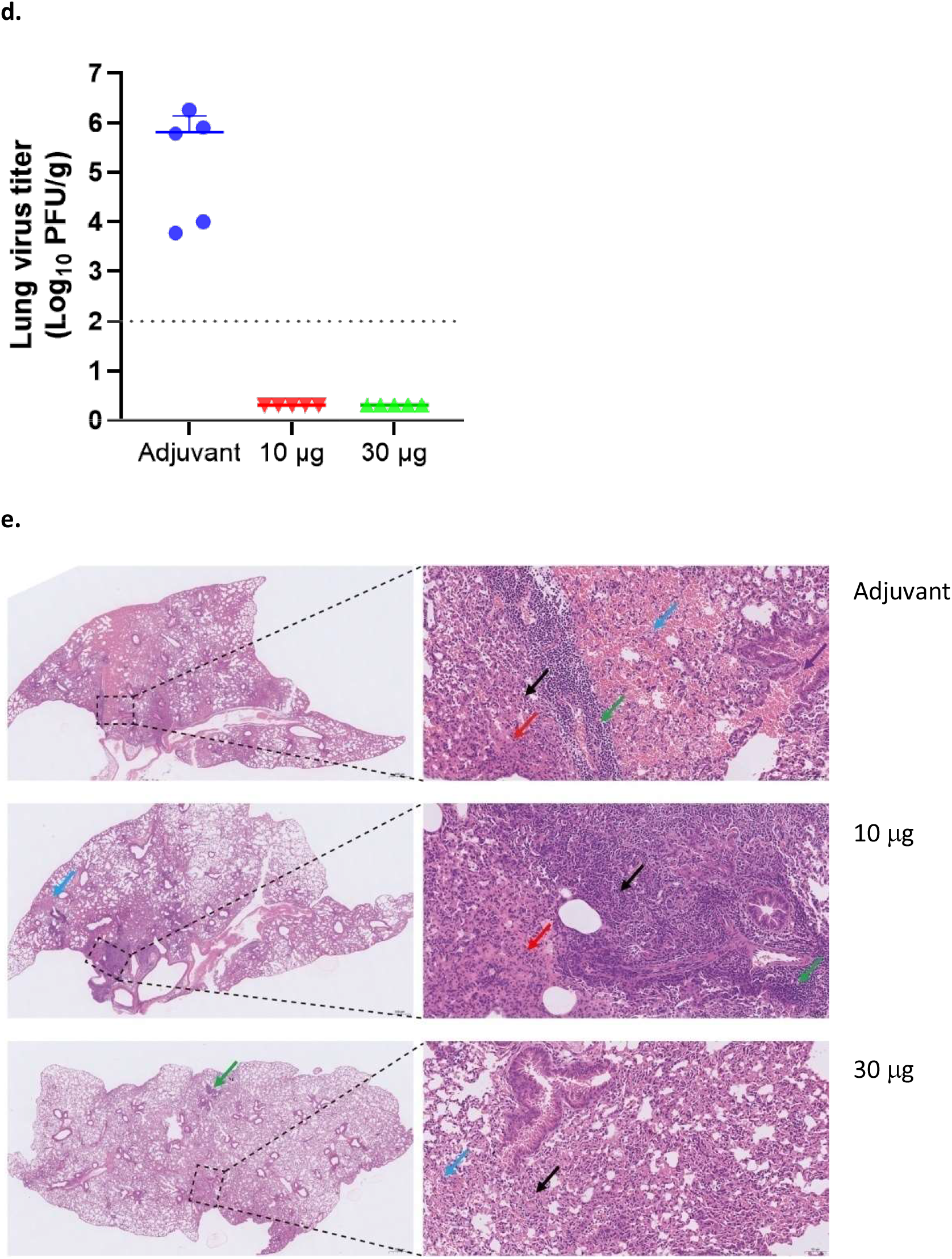
Immunogenicity and protective efficacy of PF-D-Trimer in K18-hACE2 H11transgenic mice. K18-hACE2 H11 mice were immunized with two different doses of PF-D-Trimer formulated with alum plus CpG 1018 or adjuvant only, as described in the top panel and in Material and Methods. Sera were collected and the immune responses were analyzed one week after the second injection (Day 28). Two weeks after the second injection (Day 35), the mice were exposed to SARS-CoV-2 Delta variant at 1 × 10^5^ TCID_50_ per animal by the intranasal route. **(a)** PF-D-Trimer binding antibody ELISA titers at Day 28. **(b)** Weight change of mice after exposure to SARS-CoV-2. **(c)** Survival curves of mice inoculated with SARS-CoV-2. **(d)** Viral load analysis in lung tissues at necropsy (Day 3 post-challenge) in SARS-CoV-2 infected mice. **(e)** Pathological analysis of mice lungs on Day 3 post-challenge. The image on the right is a partial enlargement of the image on the left. The black arrow represents the thickening of the alveolar wall with inflammatory cell infiltration; the blue arrow represents alveolar hemorrhage; the purple arrow represents bronchial hemorrhage; the red arrow represents pulmonary edema, and tissue exudates can be seen in the alveolar space. Dotted lines reflect assay limit of detection (100 PFU/g). Points represent individual animal, bars indicate geometric mean titers (GMT) responses (with 95% confidence intervals [CI]). • Adjuvant only, ▾PF-D-Trimer 10 µg and adjuvant, ▴PF-D-Trimer 30 µg and adjuvant. **** p≤0.0001. ANOVA F = 113.7.

The results of the lung pathological analysis at D3 after infection (Fig. 5E) were in agreement with the viral loads; the bronchial and alveolar structures of the lung tissue of the mice in the 10 μg and 30 μg adjuvanted PF-D-Trimer groups were intact and clear. There was some local alveolar wall thickening, occasionally accompanied by a small amount of inflammatory cells, lymphocyte infiltration and a small amount of angioedema, but the overall lesions were mild or moderate. However, tissue from mice of the adjuvant only group showed severe pathological features; there were large areas of alveolar wall thickening, accompanied by more inflammatory cell infiltration, more pulmonary edema and local hemorrhage at the tissue edge.

## Discussion

The PF-D-Trimer vaccine candidate comes in the form of the ectodomain of the S protein of SARS-CoV-2 fused to TD, our proprietary trimerization domain. PF-D-Trimer is in a disulfide bond-linked homotrimeric form, which permit the acquisition of important epitopes only found in the native trimeric form of the S proteins of coronaviruses and are so trimer restricted ^7^. Additionally, the trimeric form of the SARS-CoV-2 S protein adopts an open-trimer conformation potentially allowing the development of broadly protective or pan-coronavirus vaccines by exposing highly conserved region of the protein ^8^.

PF-D-Trimer is expressed at high levels in CHO cells under bioreactor culture conditions commonly found in the biopharmaceutical industry and does not require specialized equipment beyond what is readily available from recognized manufacturers in the industry. The presence of the TD domain and the α−TD domain monoclonal antibody allow an easy one-step immunoaffinity purification of the protein, which coupled with an expression titer of approximately 500 mg/L in a bioreactor, permit the implementation of a commercially favorable downstream processing, manufacturability and production.

The immunogenicity studies have shown that PF-D-Trimer adjuvanted with CpG 1018 plus alum was effective in inducing a potent immune response in four animal models: C57BL/6 mice, Sprague-Dawley rats, Syrian golden hamsters and K18-hACE2 mice, the latter being completely protected in a challenge with a live virus.

In C57BL/6 mice, vaccination with PF-D-Trimer adjuvanted with CpG 1018 and alum elicited a robust humoral immune response regardless of the dose used for immunization. We have also confirmed that a booster dose increases the antibody titer, which is very relevant since homologous and heterologous booster doses are now an integral part of anti-COVID-19 vaccine strategies ^18,19^. In addition to the humoral immune response, type 1 S-specific T-cell biased immunity likely contributes to the protection of COVID-19 convalescents ^20^. Our *in vitro* T cell analyzes demonstrated that PF-D-Trimer vaccination adjuvanted with CpG 1018 and alum elicited robust anti-recombinant SARS-CoV-2 S protein trimer specific T cells, directing the cell-mediated response towards Th1 response that may contribute to protection against SARS-CoV-2. These results are consistent with the high correlation seen between humoral and cellular immune responses to SARS-CoV-2 ^21^.

The neutralization studies have shown that SD rats immunized with PF-D-Trimer adjuvanted with alum plus CpG 1018 develop robust neutralizing antibodies against the Delta, Omicron BA.2.2 and BA.5.2 variants. We observe a reduction for the Omicron variants compare to Delta. However the reduction is less pronounced for BA.5.2 than for BA.2.2.

In our immunogenicity studies of PF-D-Trimer in the hamster model, we assessed the impact of the formulation on general antibody titers. Similar to the results observed in mice, the formulation combining PF-D-Trimer with alum plus CpG 1018 appears to generate higher titers. We have also observed that this formulation directs the immune response towards a Th1 phenotype. The hamsters’ neutralization studies have also shown that the animals immunized with the same formulation develop neutralizing antibodies directed against WA1, Delta, Omicron BA.1, BA2.12.1 and BA.4/5.

We have also demonstrated that PF-D-Trimer adjuvanted with CpG 1018 and alum induces a protective immunity in K18-hACE2 transgenic mice at the doses used for immunization. Although vaccinated mice lose weight early in the infection, they quickly return to their original weight whereas unvaccinated mice continue to lose weight until they die. Among the vaccinated mice, the viral loads were undetectable and the lungs lesions were mild or moderate, whereas the load of the control group was very high with severe pathological features. Published research results have indicated that high viral loads are associated with mortality for symptomatic and hospitalized patients who tested positive for SARS-CoV-2 ^22^.

New SARS-CoV-2 variants are emerging as the virus evolves and escapes from antibody selection pressure. In addition, recombination is extremely common in the evolutionary history of SARS-like coronaviruses and is expected to shape SARS-CoV-2 in the coming years^23^. As the Delta variant began to progressively disappear near end of 2021, a new variant, Omicron BA.1 appeared and spread very quickly. SARS-CoV-2 Delta and Omicron variants have co-circulated in the United States, allowing co-infections and possible recombination events ^11^. Studies have revealed that Omicron BA.1 was less likely to be neutralized by convalescent plasma from COVID-19 patients or plasma from people who received vaccines based on WA1 ^24, 25, 13^. Our neutralization results present a differentiating factor since PF-D-Trimer is based on the Delta strain while most of the published studies use vaccines based on WA1. We do observe a 4.7 reduction for the hamster model in the neutralization titer for Omicron BA.1, however this seems to be located in the lower region of the range obtained with the WA1 based vaccines, which varies between 6 and 23 times.

The spike protein consists of two subunits, S1 and S2. S1 mediates binding to the receptor by interaction of the receptor binding domain (RBD) with ACE2. Studies have indicated that most potent neutralizing antibodies target RBD to block the interaction with ACE2 ^26-28^. The Delta variant shares a common mutation, T478K, with BA.1 in RBD, which could partly explain the lower reduction of neutralizing antibodies when using PF-D-Trimer instead of the other vaccines based on the original WA1 spike protein. A reduction in the neutralization titers was also observed in our hamster model, but to a lesser extent, towards the original WA1 strain. The reduction can be explained by the T478K mutation but also by L452R, the latter being known to be an escape mutation for Bamlanivimab^29^.

SARS-CoV-2 is in constant evolution. BA.2.12.1 evolved from BA.2 in the United State in the spring of 2022 while Omicron BA.4 and BA.5 were identified in South Africa in January and February 2022. These variants are outcompeting the previously circulating BA.1 and BA.2 variants. In the summer of 2022, BA.5 was responsible for almost 90% of COVID-19 cases in the United States. Meanwhile, BA.4 was responsible for about 5% of cases ^30^. The spike protein of BA.2.12.1 contains the L452Q and S704L mutations in addition to the known mutations in BA.2. The spike proteins of BA.4 and BA.5 are identical, each with the additional mutations Δ69-70, L452R, F486V. Studies ^31,32^, identified that the specific mutations found in BA.2.12.1 and BA.4/5 contribute to antibody resistance, BA.4/5 showing a more pronounced reduced neutralization than BA.2.12.1 to the vaccination induced by the Pfizer– BioNTech BNT162b2 vaccine. Hachmann, N.P. et al. ^33^ have shown that the BA.2.12.1, BA.4, and BA.5 subvariants substantially escape neutralizing antibodies in sera from participants who had been infected with BA.1 or BA.2.

As the newly emerged variants represent a significant proportion of new cases of infection, it becomes imperative to develop a vaccine that can generate adequate protection against them. Interestingly, Qu, P. et al. ^13^ demonstrates that sera from unvaccinated or vaccinated Delta-wave ICU patients more strongly neutralize BA.4/5 and BA.2.12.1 compared to BA.1 and BA.2, while infection during the BA.1 wave did not appear to offer effective protection against the new sublineages. We also observe higher neutralization titers against BA.5.2 compared to BA.2.2 with sera from SD rats immunized with PF-D-Trimer. It is worth noting that the BA.4/5 sublineages and Delta share the immune escape associated L452R mutation ^34^, which could explain why PF-D-Trimmer can induce high neutralization titers against BA.5.2.

It is known that neutralization antibody levels are highly predictive of immune protection ^35^. Our results are showing that adjuvanted PF-D-Trimer induces high neutralization titers for different SARS-CoV-2 variants including Omicron BA.2.2 and BA.5.2. In sum, the Delta spike protein could represent an effective booster vaccine antigen by activating cross-neutralizing B cells and a T cell response.

Delta infection was found to be broadly immunogenic in mice. Sera from likely Delta breakthrough cases can also neutralize VLPs generated with S genes from the Delta and BA.1 strains ^36^. In SARS-CoV-2 pseudotyped virus neutralization assays, a Delta breakthrough infection (BTI) in double vaccinated subjects induces more potent cross-neutralizing antibodies against variants including BA.1 than Delta infection in unvaccinated subjects. As such a Delta BTI was determined to be an effective booster which could provide broad protection ^37^. In vaccinated Delta-infected patients, RBD IgG titers are superior compared to BA.1-infected subjects and the CD8+ T cells are more spike oriented ^38^.

PF-D-Trimer consists of the full Delta S-protein ectodomain including the S1 and S2 subdomains, trimerized by the Trimer Domain. Following immunization, the immune response appears to target more variable and likely more immunogenic epitopes in the RBD and N-terminal domain of S1 of SARS-CoV-2 strains ^39,40^. The S2 subunit, which is more sequence conserved than S1, harbors a significant proportion of antibody-targeted epitopes in the response to SARS-CoV-2 infection or vaccination. Thus, cross-reacting antibodies targeting conserved regions in SARS-CoV-2 S2, such as the fusion peptide, heptad repeats, stem helix, or membrane proximal regions, are also stimulated by SARS-CoV-2 infection or vaccination ^41,42^. One such antibody is HCLC-031 that efficiently neutralizes Delta and Omicron BA.1 ^43^. The oligomerization process modulates the antigenicity of the coronavirus S protein, in addition to the epitopes located on the monomeric form, some are trimer restricted ^7^. Also, the trimeric form of SARS-CoV-2 S adopts an open conformation that exposes the S2 interface of the trimer and its highly conserved epitopes ^8^. It is thus likely that PF-D-Trimer is a more complete immunogen than vaccines based on domains or subdomains of the S protein.

Our data demonstrate that PF-D-Trimer adjuvanted with alum plus CpG 1018 can induce cross-reactive humoral and cellular immune responses in various animal species and protective immunity against a homologous SARS-CoV-2 challenge in K18-hACE2 H11 transgenic mice. The adjuvanted PF-D-Trimer vaccine candidate has already passed the necessary safety assessments for advancement to human clinical trials as well as Investigator Initiated Trials (IITs) to define the clinical objectives. Together, the results support the advancement of adjuvanted PF-D-Trimer into human clinical studies to further demonstrate safety, immunogenicity and vaccine efficacy. The advantage brought by the Trimer Domain for the stabilization of the antigen, the streamlined downstream processing and CMC makes it possible to envisage a rapid scale-up of the production and rapid antigen interchangeability during the inevitable evolution of SARS-CoV-2.

## Author contributions

Z.Z. carried out the stable CHO cell clone selections and characterization as well as the recombinant protein functional assays. J.Z. was in charge of the bioprocess development for protein expression in bioreactors. P.N. was in charge of the adjuvant development and immunoassays. B.H. designed and performed the neutralization assays with live SARS-CoV-2 virus in BL-3 facility. S.D. performed the protein purification and characterization. Q.X. performed the antibody production and purification. Q.H. performed the immunoassays (ELISA and ELISpot). G.L. performed the plasmids construction and protein expression. Y. Xia. performed the animal immunizations and assays. M.L. performed the plasmids construction. C.W. performed the adjuvant development and immunoassays. Z-R.Z. performed the hACE2 mice live SARS-CoV-2 challenges. B.Z. supervised the hACE2 mice live SARS-CoV-2 challenges. K.C. supervised the neutralization assays with live SARS-CoV-2 virus. N.X. was the Q&A supervisor. Z.F. supervised the project. N.J. wrote the manuscript with support from Y. Xu, conceived the original idea and managed the project. Y. Xu. conceived the original idea, managed and supervised the project. B.L. managed and supervised the project.

## Conflicts of interest

J.Z., P.N., Q.X., C.W. N.X., Z.F. and N.J. are employees in Wuhan Binhui Biopharmaceutical Co., Ltd.

S.D., G.L. and Y. Xia are employees in Nova Biologiques Inc.

Y. Xu owns shares in Nova Biologiques Inc.

B.L. owns shares in Wuhan Binhui Biopharmaceutical Co., Ltd.

The remaining authors have no conflicts of interest to declare.

## Materials & Correspondence

Correspondence and requests for materials should be addressed to B.L. or to Y.X. or to B.Z. or K.C.

## Data Availability Statement

The datasets generated during and/or analysed during the current study are available from the corresponding authors on reasonable request.

## Materials and Methods

### Animals and biosafety experiments

All the animals’ experiments, except the experiments with the K18-hACE2 H1 mice, were approved by the Hubei University of Technology Laboratory Animal Ethics Review Committee (Approval number: 2021018). The K18-hACE2 H1 mice viral infections were conducted in the biosafety level 3 (BSL-3) facility at Wuhan Institute of Virology under a protocol approved by the Laboratory Animal Ethics Committee of Wuhan Institute of Virology, Chinese Academy of Sciences (Permit number: WIVA26202201). All works with live SARS-CoV-2 virus titration and neutralization assays were performed inside biosafety cabinets in the biosafety level 3 (BSL-3) facility at Hubei Provincial Center for Disease Control and Prevention. Specific pathogen-free (SPF) female C57BL/6 mice (6-8 week old), SPF female Syrian hamsters (5-6 week old) and 6 week old Sprague–Dawley rats were purchased from the Hubei Laboratory Research Center. H11-K18-hACE2 male mice (6-8 week) were purchased from Jiangsu Jicui Yaokang Biotechnology Co., Ltd. Animals had free access to water and food in a controlled environment with a 12-hour light/dark cycle (temperature: 16–26 °C, humidity: 40–70%).

### Protein trimerization, expression and purification

The SARS-CoV-2 S glycoprotein B.1.617.2 sequence was downloaded from GISAID (accession number EPI_ISL_1970349). The S gene was codon optimized for high level expression in CHO mammalian cells and biochemically synthesized by Genscript (China). Compared to the reference gene, the following modifications have been made during the synthesis: three mutations (RRAR to GSAS) were introduced in the S1/S2 furin site of the full-length SARS-CoV-2 S protein along with the 2P mutation; the transmembrane and the cytoplasmic domains were deleted; and the trimerization domain (TD) was added at the C-terminus. The recombinant SARS-CoV-2 S gene was cloned in the pGenHT1.0-DGV plasmid with a 5’CMV promoter and a 3’ polyA sequence. The gene sequence of the recombinant SARS-CoV-2 S protein with GSAS, 2P and TD was confirmed by DNA sequencing. The linearized target plasmid was transfected into CHOK1-GenS cells to generate a stable cell pool. Subsequent screening for high-titer production clones, process optimization and a fed-batch serum free cell culture process in bioreactor, led to the production of recombinant PF-D-Trimer as a highly secreted protein in the culture medium of CHO cells.

The α−TD monoclonal antibody was produced in Sf9 insect cells by using a Bac-to-Bac recombinant baculovirus system (Invitrogen). The monoclonal antibody manufacturing process started with the revival, expansion and production of the Sf9 cells from the working cell bank (WCB) into shake flasks and bioreactor using serum-free medium (Vbiosci). The cultured cells were infected by inoculation of a recombinant baculovirus carrying the α−TD monoclonal antibody genes. At the end of the infection phase, the cell culture supernatant with the secreted antibody was harvested by centrifugation. The clarified medium was subjected to a purification process by MabSelect PrismA (Cytiva) affinity chromatography according to the manufacturer’s instruction. The affinity chromatography eluate was subjected to concentration and diafiltration using TFF (Sartorius Stedim Biotech GmbH). In order to produce the immunoaffinity resin for PF-D-Trimer purification, the α−TD monoclonal antibody was coupled to NHS-activated Sepharose 4 Fast Flow (Cytiva) according to the manufacturer’s instructions.

The PF-D-Trimer active substance manufacturing process started with the revival, expansion and production of the CHO cells from the working cell bank (WCB) into shake flasks and bioreactor (Applikon) using serum-free medium (Gibco). At the end of the growth phase, the cell culture supernatant was harvested by depth filtration (Sartorius Stedim Biotech GmbH). The clarified medium was subjected to a purification process by α−TD monoclonal antibody immunoaffinity chromatography to capture PF-D-Trimer, and virus inactivation by a low pH treatment. The immunoaffinity chromatography eluate was subjected to concentration and diafiltration using tangential flow filtration (TFF) (Sartorius Stedim Biotech GmbH) and final 20 nm nanofiltration (Pall Corporation) to obtain a purified SARS-CoV-2 recombinant PF-D-Trimer active substance.

### PF-D-Trimer purity, size, aggregation and fragmentation analysis by SEC-HPLC

The α−TD immunoaffinity purified PF-D-Trimer was analyzed by Size-Exclusion Chromatography (SEC-HPLC) using a Shimadzu LC-2030 HPLC (Shimadzu Corporation, Japan) with an analytical SRT SEC-300 7.8×300mm column (Sepax). Phosphate-buffered saline (PBS) was used as the mobile phase with OD_280_ nm detection over a 20-min period at a flow rate of 1 ml/min.

### Negative stain electronic microscopy

To characterize the oligomerization status of immunoaffinity purified PF-D-Trimer, 20 µl of samples were added dropwise to 200-mesh grids (Fanghua Film Copper Mesh) and incubated at room temperature for 10 minutes. Then the grids were negatively stained with 2% phosphotungstic acid for 3 minutes, and the remaining liquid was removed with a filter paper. The prepared samples were observed with a HT7800 (Hitachi) transmission electron microscope.

### Immunogenicity of PF-D-Trimer

Twenty-eight C57BL/6 mice were randomly divided into 4 immunization groups; PBS with 50 μg aluminum hydroxide (hereafter abbreviated as alum) plus 10 μg CpG 1018 (General Biol), 5 μg PF-D-Trimer with 50 μg alum plus 10 μg CpG 1018, 10 μg PF-D-Trimer with 50 μg alum plus 10μg CpG 1018, 20 μg PF-D-Trimer with 50 μg alum plus 10 μg CpG 1018. The total injection volume of the formulated vaccines was 100 µL per dose. Three doses were administered subcutaneously at Day 0, Day 14 and Day 28. The sera of immunized mice were collected by retro-orbital bleeding at Day 21 and Day 35 to detect the SARS-CoV-2-specific IgG endpoint GMT. Spleens were removed for ELISpot assays after sacrifice by cervical dislocation at Day 51.

Sprague-Dawley rats were immunized intramuscularly (IM) with two different doses of PF-D-Trimer adjuvanted with 375 µg alum plus 0.75 mg CpG 1018: five rats with 10 µg of protein, and six rats with 30 µg of protein. The total injection volume of the formulated vaccines was 500 µL per IM dose. Three IM doses were administered (at Day 0, Day 22 and Day 43). The sera were collected at Day 22, Day 43 and Day 71 to detect SARS-CoV-2 specific neutralizing antibodies as described below.

Syrian hamsters were divided into 7 groups (PBS alone, PBS with 75 μg alum plus 150 μg CpG 1018, 10 μg of PF-D-Trimer alone, 5 μg PF-D-Trimer only with 75 μg alum, 10 μg PF-D-Trimer only with 75 μg alum, 5 μg PF-D-Trimer with 75 μg alum plus 150 μg CpG 1018, 10 μg PF-D-Trimer with 75 μg alum plus 150 μg CpG 1018). The total injection volume of the formulated vaccines was 300 µL per intra-muscular (IM) dose. Three IM injections were administered (at Day 0, Day 22 and Day 90). The sera were collected at Day 42 and Day 110 to detect the SARS-CoV-2-specific IgG endpoint GMTs and neutralizing antibodies as described below. Spleens were removed for ELISpot assays after sacrifice at Day 110.

### SARS-CoV-2-specific antibody endpoint GMT measurement with ELISA

Specific IgG antibody endpoint GMT were measured by ELISA. Briefly, sera serially diluted in ELISA buffer (1% BSA with 0,05% Tween-20) were added (100 µl/well) to 96-well plates (Costar) coated with recombinant SARS-CoV-2 S protein antigen (Novozan Biotechnology Co., Ltd.) and blocked with ELISA buffer for 120 min at 37°C. After three washes with wash buffer (PBS with 0,05% Tween-20), the plates were incubated at 37°C for 30 minutes with horseradish peroxidase-conjugated goat anti-mouse IgG (Proteintech) or goat anti-hamster IgG (Shanghai Universal Biotech Co., Ltd.), both diluted in ELISA buffer at a 1 in 10000 dilution. The plates were then washed 3 times with wash buffer. Signals were developed using TMB substrate (Solarbio). The colorimetric reaction was stopped by the addition of 2M H_2_SO_4_. Finally, the absorbance (450 nm) was measured with a Varioskan LUX Multimode Microplate Reader (Thermo). The IgG endpoint GMT were defined as the dilution fold with an optical density exceeding the average background plus 3 times the standard deviation (sera from the PBS control groups).

### ELISpot assays

Spleens from immunized animals were removed at Day 51 for the mice and at Day 110 for the hamsters and splenocytes were isolated for protein S-specific T-cell detection with the ELISpotPLUS mouse IFN-γ or ELISpotPLUS hamster IFN-γ kits (Mabtech) according to the manufacturer’s instructions. Briefly, 3∼5×10^5^ splenocytes/well and 4 μg/well SARS-CoV-2 spike protein B.1.617.2 (Nanjing Vazyme Biotech Co., Ltd) were mixed and incubated at 37°C. After 48 hours followed by 5 washes with PBS, the probe antibody was added at a concentration of 1 μg/μl and incubated at room temperature for 2 hours. After washing 5 times with PBS, alkaline phosphatase labeled streptavidin was added at a dilution of 1 in 1000 and incubated for 1 hour at room temperature. After washing 5 times with PBS, alkaline phosphatase labeled streptavidin was added at a dilution of 1 in 1000 and incubated for 1 hour at room temperature. After washing 5 times with PBS, the chromogenic solution (BCIP/NBT-plus) was added and incubated at room temperature for 10 minutes. Color development was then stopped with deionized water. The number of dots in the wells of the ELISpot plate was analyzed using an ELISpot reader system (AID ELISpot Reader Classic and AID ELISpot Reader Software, Autoimmun Diagnostika GmbH).

### Pseudovirus-based neutralization assay

The 50% neutralization titer (NT50) was measured using a modification of the procedure from Nie, J. *et al*. ^44^. Briefly, the sera from immunized animals were diluted 3 times, starting from 1:33.33, and incubated with SARS-CoV-2 pseudovirus (2×10^4^ TCID_50_/mL) at 37°C for 60 min. DMEM without serum was used as a negative control group. Then the HEK293T-hACE2 cells were added to each well (2×10^4^ cells/well) and incubated at 37°C for 48 hours. Luciferase activity, which reflects the degree of SARS-CoV-2 pseudovirus transduction, was measured using Bio-Lite Luciferase Assay System (Vazyme). The NT50, calculated by the Reed-Muench method ^45^, was defined as the fold dilution which obtained more than 50% inhibition of pseudovirus transduction compared to the control group.

### SARS-CoV-2 neutralization assays

Vero E6 cells (2.5 × 10^4^ cells/well) were seeded in 96-well plates and incubated overnight. Sera were inactivated at 56 °C for 30 min and diluted in serum-free DMEM at an initial dilution factor of 8, and then further serially diluted. The diluted sera were then mixed in a 1:1 ratio with the SARS-CoV-2 viruses (Hubei Provincial Centre for Disease Control and Prevention, WA1-Hu-1, Delta YJ20210701-01, Omicron BA.1 249099, Omicron BA.2.2 YJ20220413-11 and Omicron BA.5.2 YJ20220704-03 at 100 TCID_50_/100 μl and incubated one hour at 37°C. Then, the diluted sera/virus mixtures were added to the Vero cells and incubated at 37°C with 5% CO_2_ for four days. The cells were monitored for cytopathic effect (CPE) every 24 hours under an inverted microscope for each sera dilution. The neutralization end point 50%, the dilution of serum that can protect 50% of cells from CPE, was calculated by the Reed-Muench method ^45^ to obtain the neutralizing antibody titer of each serum.

### K18-hACE2 H11 mice immunization and challenge study

K18-hACE2 H11 (C57BL/6JGpt) male transgenic mice were divided in 5 vaccination groups (#1 not immunized, 6 animals; #2 PBS control group, 7 animals; #3 PBS with 125 µg alum plus 750 µg CpG, 10 animals; #4 10 µg PF-D-Trimer with 125 µg alum plus 750 µg CpG, 10 animals; #5 30 µg PF-D-Trimer with 125 µg alum plus 750 µg CpG, 10 animals). Blood was collected before the immunization as a pre-immune control. Two subcutaneous 500 µl (5 dorsal multipoint injections of 100 µl) doses were administered (at Day 0 and Day 21). The sera were collected at Day 28 to detect the SARS-CoV-2-specific IgG endpoint GMT and neutralizing antibodies as described. At Day 35, mice from groups #3 to #5 were transferred to a BSL3 laboratory (Wuhan Institute of Virology). For the challenge study, the mice were intranasally infected with SARS-CoV-2 Delta (CRST: 1633.06.IVCAS 6.7593) at 1 × 10^5^ plaque-forming unit (PFU) (50 μl per animal). Change in body weight and determination of survival rate were monitored daily. Half of the animals were euthanized at 3 days post infection (dpi) and the remaining animals in each group were followed until day 7. Lung tissue samples from the euthanized mice were collected to determine the virus titer by plaque assay or fixed with 4% paraformaldehyde and stained with hematoxylin and eosin for pathological analysis. Quantification of viral titer in lung tissue by cell culture infectious assay (TCID_50_) was performed as follows. Vero E6 cells were plated into 24-well plates 1 day in advance at a density of 10^5^ cells/well. Lung homogenates were prepared from 100 mg of lung tissue in 1 ml PBS. The next day, the cells were infected by tenfold serial dilutions of 100 μl lung homogenate supernatant for each sample and incubated at 37°C for 1 hour. The virus dilution was then removed and 1% methylcellulose was added. The plates were incubated for 4 days at 37°C. The upper layer in the wells was discarded and 1 ml of fixative staining solution (3.7% formaldehyde + 1% crystal violet) was added. The cells were treated overnight at room temperature. After the fixative staining solution was rinsed with running water and dried, the number of viral plaques was counted. The plaque forming units per mL (PFU/ml) were determined using the following formula: (# plaques × dilution factor)/0.1 ml. An absence of plaque was scored as <1 and used to calculate the lower limit of detection. The PFU/ml values were adjusted for a volume of 1 ml and 1 g of tissue to calculate the plaque forming units per gram of tissue.

### Statistical Analysis

Data were arranged in Excel and analyzed using GraphPad Prism 8.0.1 or 8.0.2. Two-tailed t-tests were used to compare the means of each two PF-D-Trimer immunized groups. Ordinary one-way ANOVA with Dunnett’s multiple comparison was used to analyze the differences among groups.

## References

1 WHO. WHO Coronavirus disease (COVID-19) Weekly Epidemiological Update and Weekly Operational Update, <https://www.who.int/emergencies/diseases/novel-coronavirus-2019/situation-reports> (2022).

2 Zhou, P. et al. A pneumonia outbreak associated with a new coronavirus of probable bat origin. Nature, 7798 (2020).

3 Wrapp, D., Wang, N., Corbett, K. S., Goldsmith, J. A. & Mclellan, J. S. Cryo-EM Structure of the 2019-nCoV Spike in the Prefusion Conformation. Science 1260– 1263 (2020).

4 Abdulla, Z. A., Al-Bashir, S. M., Al-Salih, N. S., Aldamen, A. A. & Abdulazeez, M. Z. A Summary of the SARS-CoV-2 Vaccines and Technologies Available or under Development. Pathogens 10, 788 (2021).

5 Dai, L. & Gao, G. Viral targets for vaccines against COVID-19. Nature reviews. Immunology 21, 73–82, doi:10.1038/s41577-020-00480-0 (2021).

6 Chen, W. H., Strych, U., Hotez, P. J. & Bottazzi, M. E. The SARS-CoV-2 Vaccine Pipeline: an Overview. Current Tropical Medicine Reports, 61–64 (2020).

7 Delmas, B. & Laude, H. Assembly of coronavirus spike protein into trimers and its role in epitope expression. Journal of Virology 64, 5367–5375 (1990).

8 Costello, S. M., Shoemaker, S. R. & Hobbs, H. T. The SARS-CoV-2 spike reversibly samples an open-trimer conformation exposing novel epitopes. Nature Structural & Molecular Biology, 229–238 (2022).

9 Yaniv, K. et al. Managing an evolving pandemic: Cryptic circulation of the Delta variant during the Omicron rise. Science of The Total Environment 836, 155599, doi:https://doi.org/10.1016/j.scitotenv.2022.155599 (2022).

10 Zou, J., Xie, X., Liu, M., Shi, P.-Y. & Ren, P. SARS-CoV-2 Delta breakthrough infections in vaccinated patients. bioRxiv : the preprint server for biology, doi:10.1101/2022.04.12.488092 (2022).

11 Bolze, A. et al. Evidence for SARS-CoV-2 Delta and Omicron co-infections and recombination. medRxiv, doi:10.1101/2022.03.09.22272113 (2022).

12 Wawina-Bokalanga, T. et al. Genomic evidence of co-identification with Omicron and Delta SARS-CoV-2 variants: a report of two cases. Int J Infect Dis 122, 212–214, doi:10.1016/j.ijid.2022.05.043 (2022).

13 Qu, P. et al. Neutralization of the SARS-CoV-2 Omicron BA.4/5 and BA.2.12.1 Subvariants. The New England journal of medicine 386, 2526–2528, doi:10.1056/NEJMc2206725 (2022).

14 Pallesen, J., Wang, N., Corbett, K. S., Wrapp, D. & Mclellan, J. S. Immunogenicity and structures of a rationally designed prefusion MERS-CoV spike antigen. Proc Natl Acad Sci U S A 114, E7348 (2017).

15 Xu Yan, J. N., Li Gai, Xia Yan, Deng Shuang. Heterotrimerization domain, heterotrimerization fusion protein, preparation method and application. China patent CN113480616A (2021).

16 Rössler, A., Riepler, L., Bante, D., von Laer, D. & Kimpel, J. SARS-CoV-2 Omicron Variant Neutralization in Serum from Vaccinated and Convalescent Persons. The New England journal of medicine 386, 698–700, doi:10.1056/NEJMc2119236 (2022).

17 Yinda, C. et al. K18-hACE2 mice develop respiratory disease resembling severe COVID-19. PLoS Pathogens, doi:10.1371/journal.ppat.1009195 (2021).

18 Munro, A. et al. Safety and immunogenicity of seven COVID-19 vaccines as a third dose (booster) following two doses of ChAdOx1 nCov-19 or BNT162b2 in the UK (COV-BOOST): a blinded, multicentre, randomised, controlled, phase 2 trial. Lancet (London, England) 398, 2258–2276, doi:10.1016/s0140-6736(21)02717-3 (2021).

19 NCIRD. National Center for Immunization and Respiratory Diseases (NCIRD), Division of Viral Diseases. Update, <https://www.cdc.gov/coronavirus/2019-ncov/vaccines/booster-shot.html> (2022).

20 Weiskopf, D. et al. Phenotype and kinetics of SARS-CoV-2-specific T cells in COVID-19 patients with acute respiratory distress syndrome. Science immunology 5, doi:10.1126/sciimmunol.abd2071 (2020).

21 Österdahl, M. F. et al. Concordance of B and T cell responses to SARS-CoV-2 infection, irrespective of symptoms suggestive of COVID-19. J Med Virol, doi:10.1002/jmv.28016 (2022).

22 Pujadas, E. et al. SARS-CoV-2 viral load predicts COVID-19 mortality. The Lancet. Respiratory medicine, 9, doi:10.1016/s2213-2600(20)30354-4 (2020).

23 Müller, N. F., Kistler, K. E. & Bedford, T. A Bayesian approach to infer recombination patterns in coronaviruses. Nat Commun 13, 4186, doi:10.1038/s41467-022-31749-8 (2022).

24 Planas, D. et al. Considerable escape of SARS-CoV-2 Omicron to antibody neutralization. Nature 602, 671–675, doi:10.1038/s41586-021-04389-z (2022).

25 Dejnirattisai, W. et al. Omicron-B.1.1.529 leads to widespread escape from neutralizing antibody responses. bioRxiv : the preprint server for biology, doi:10.1101/2021.12.03.471045 (2021).

26 Dejnirattisai, W., Zhou, D., Ginn, H. M., Duyvesteyn, H. & Screaton, G. R. The antigenic anatomy of SARS-CoV-2 receptor binding domain. Cell, 2183–2200 (2021).

27 Lan, J. et al. Structure of the SARS-CoV-2 spike receptor-binding domain bound to the ACE2 receptor. Nature, 215–220 (2020).

28 Liu, C. et al. Reduced neutralization of SARS-CoV-2 B.1.617 by vaccine and convalescent serum. Cell 184, 4220-4236.e4213, doi:10.1016/j.cell.2021.06.020 (2021).

29 Starr, T., Greaney, A., Dingens, A. & Bloom, J. Complete map of SARS-CoV-2 RBD mutations that escape the monoclonal antibody LY-CoV555 and its cocktail with LY-CoV016. Cell reports. Medicine 2, 100255, doi:10.1016/j.xcrm.2021.100255 (2021).

30 CDC. COVID Data Tracker, <https://covid.cdc.gov/covid-data-tracker> (2022).

31 Wang, Q. et al. Antibody evasion by SARS-CoV-2 Omicron subvariants BA.2.12.1, BA.4, & BA.5. Nature, doi:10.1038/s41586-022-05053-w (2022).

32 Arora, P. et al. Augmented neutralisation resistance of emerging omicron subvariants BA.2.12.1, BA.4, and BA.5. The Lancet. Infectious diseases, doi:10.1016/s1473-3099(22)00422-4 (2022).

33 Hachmann, N. et al. Neutralization Escape by SARS-CoV-2 Omicron Subvariants BA.2.12.1, BA.4, and BA.5. The New England journal of medicine 387, 86–88, doi:10.1056/NEJMc2206576 (2022).

34 Motozono, C. et al. SARS-CoV-2 spike L452R variant evades cellular immunity and increases infectivity. Cell Host & Microbe 29, 1124-1136.e1111, doi:https://doi.org/10.1016/j.chom.2021.06.006 (2021).

35 Khoury, D. S. et al. Neutralizing antibody levels are highly predictive of immune protection from symptomatic SARS-CoV-2 infection. Nature Medicine 27, 1205–1211, doi:10.1038/s41591-021-01377-8 (2021).

36 Suryawanshi, R. et al. Limited Cross-Variant Immunity after Infection with the SARS-CoV-2 Omicron Variant Without Vaccination. medRxiv : the preprint server for health sciences, doi:10.1101/2022.01.13.22269243 (2022).

37 Lechmere, T. et al. Broad Neutralization of SARS-CoV-2 Variants, Including Omicron, following Breakthrough Infection with Delta in COVID-19-Vaccinated Individuals. mBio, 2, doi:10.1128/mbio.03798-21 (2022).

38 Kared, H. et al. Immunity in Omicron SARS-CoV-2 breakthrough COVID-19 in vaccinated adults. medRxiv, 22269213, doi:10.1101/2022.01.13.22269213 (2022).

39 Amanat, F. et al. SARS-CoV-2 mRNA vaccination induces functionally diverse antibodies to NTD, RBD, and S2. Cell 184, 3936-3948.e3910, doi:10.1016/j.cell.2021.06.005 (2021).

40 Piccoli, L. et al. Mapping Neutralizing and Immunodominant Sites on the SARS-CoV-2 Spike Receptor-Binding Domain by Structure-Guided High-Resolution Serology. Cell 183, 1024-1042.e1021, doi:10.1016/j.cell.2020.09.037 (2020).

41 Voss, W. N. et al. Prevalent, protective, and convergent IgG recognition of SARS-CoV-2 non-RBD spike epitopes. Science 372, 1108–1112, doi:10.1126/science.abg5268 (2021).

42 Ladner, J. T. et al. Epitope-resolved profiling of the SARS-CoV-2 antibody response identifies cross-reactivity with endemic human coronaviruses. Cell Rep Med 2, 100189, doi:10.1016/j.xcrm.2020.100189 (2021).

43 Hu, J. et al. A spike protein S2 antibody efficiently neutralizes the Omicron variant. Cellular & Molecular Immunology 19, 644–646, doi:10.1038/s41423-022-00847-4 (2022).

44 Nie, J. et al. Quantification of SARS-CoV-2 neutralizing antibody by a pseudotyped virus-based assay. Nat Protoc 15, 3699–3715, doi:10.1038/s41596-020-0394-5 (2020).

45 Reed, L. J. & Muench, H. A simple method of estimating fifty percent endpoints. American Journal of Epidemiology 27, 493–497, doi:10.1093/oxfordjournals.aje.a118408 (1938).

